# Learning continuous updating and control of neural population activities from sparse time samples

**DOI:** 10.64898/2026.09.28.754781

**Authors:** Zezhao Wu, Mingsha Zhang, Ning Qian

## Abstract

The brain frequently updates or controls its internal representations of sensory or motor variables over time. An example is transsaccadic updating of stimuli’s retinotopic positions: When the eye moves in one direction, the positions are updated continuously in the opposite direction, and the process is controlled by the corollary discharge (CD) of the saccade motor command. We previously trained neural networks to perform such updating using the desired time courses of stimuli’s positions at every time step. However, this may not be biologically plausible because, before a system learns to perform the task, it may not have access to complete time courses and the actual visual inputs are delayed and thus incorrect. Here we show that by adding the continuity equation to the loss, we can train neural networks with only stimuli’s pre- and post-saccadic retinotopic positions (without specifying the intermediate transitions), and with heavily down-sampled time courses. Moreover, both the connectivity patterns and the CD control signal can be learned simultaneously. Since the continuity equation can be applied to waves of neural population activities that are distributed representations of variables, our work also suggests an advantage of distributed representations which are ubiquitous in the brain.

## 1 Introduction

The brain internally represents various useful variables of the body and world for sensorimotor processing. These representations have to be updated/controlled through time with changing situations. An example is transsaccadic updating of stimuli’s retinotopic positions. Assume that the brain represents a stimulus of interest at a certain retinotopic position. If the eye saccades to the right by 10 deg, then the stimulus’ positional representation needs to be updated to the left by 10 deg. Many studies have investigated neural mechanisms of the updating process and a circuit model has been proposed recently (Duhamel, Colby, & Goldberg, 1992; Neupane, Guitton, & Pack, 2016; Sommer & Wurtz, 2002; Umeno & Goldberg, 1997; Wang et al., 2016, 2025, 2024). The model consists of a population of cells with receptive fields (RFs) covering a range of retinotopic positions and represents a stimulus’s position as a population activity bump centered at that position (Fig. 1a). The activity bump is sustained by a symmetric, center-surround connectivity among the cells (Fig. 1b, red curve). Around the time of a saccade (from about 50-100 ms before the saccade onset to about 100 ms after the saccade offset), the population activity bump is shifted backward (against the saccade) as a wave by a directional connectivity (Fig. 1b, blue curve) gated by the corollary discharge (CD) of the saccade command. The backward updating of the population activity is equivalent to the forward remapping of RFs observed in many studies (Duhamel et al., 1992; Qian, Goldberg, & Zhang, 2023; Wang et al., 2025, 2024).

**Fig. 1.**
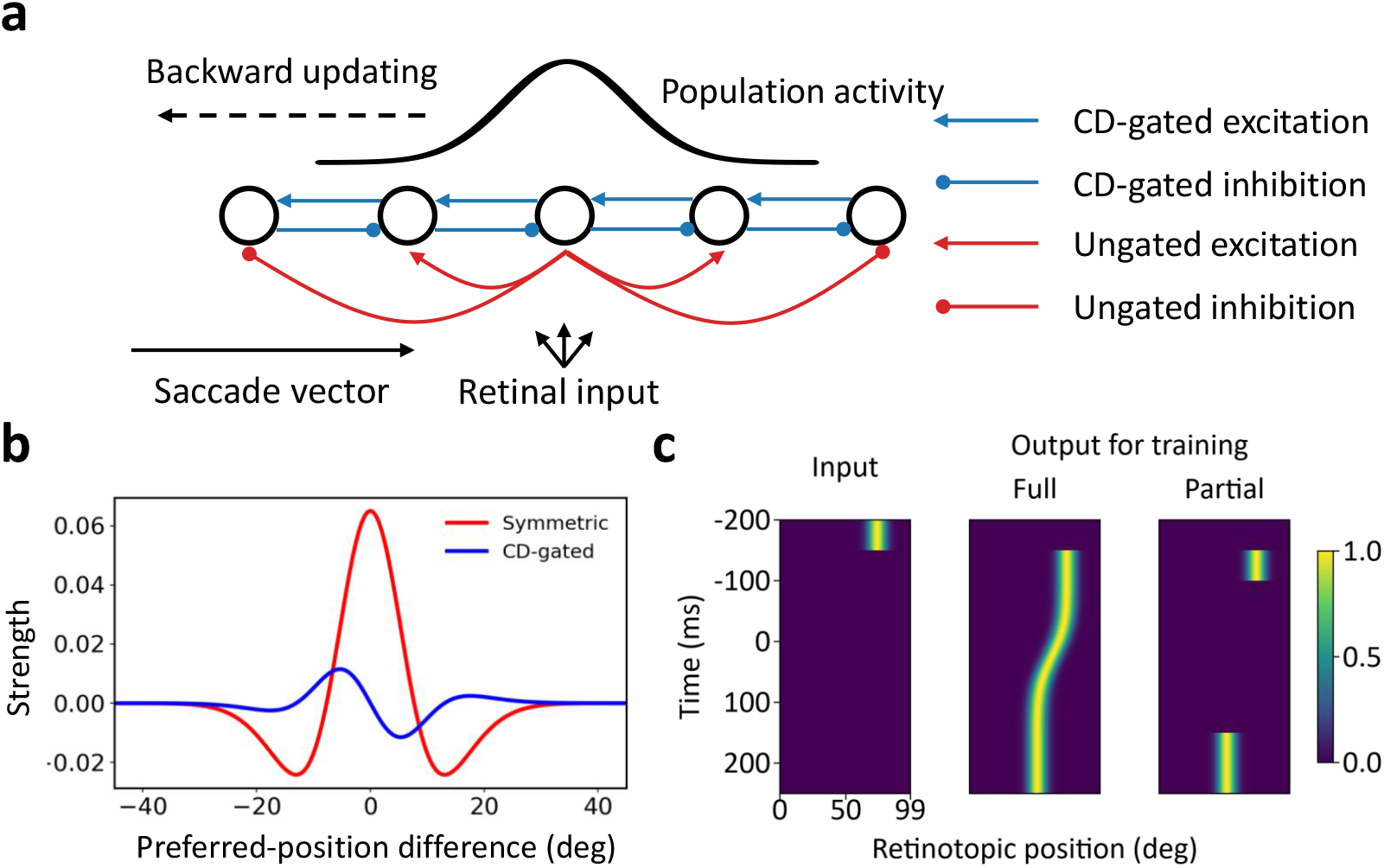
Recurrent neural networks, and the input and desired output for training. (a) Schematic of transsaccadic updating of the population activity bump. For the illustrated rightward saccade, the bump shifts in the leftward direction. The activities of the recurrent units are also the model output. Blue and red arrows denote CD-gated and ungated recurrent connections, respectively. To avoid clutter, only subsets of the connections are shown. (b) The connection strengths used in our previous models, plotted as a function of the difference between the preferred positions of the connected units. The ungated connections (red) are symmetric with central excitation and surround inhibition. The CD-gated connections (blue, for rightward saccades) follow an anti-symmetric, directional pattern. (c) Example heatmaps showing the brief visual input, the desired output in the full-output condition, and the desired output in the partial-output condition. We used different starting positions, saccade directions and amplitudes in training and testing. Time is relative to the saccade onset at 0 ms.

The circuit model is consistent with relevant physiological and psychophysical data (Wang et al., 2025, 2024) but it is less clear how the required connectivity patterns come to exist. To address the issue, we previously trained the model with random initial connections (some are gated by the CD and some are not) on the task of transsaccadic updating and found that the required connectivity patterns emerge after the training (Wang et al., 2024). However, the training used the desired time courses of stimuli’s retinotopic positions sampled at every time step. The brain may not have access to such complete trajectories before it learns the correct connectivity patterns. In particular, the updating/remapping in the lateral intraparietal area (LIP) and frontal eye fields (FEF) starts before the saccade onset, and this predictive aspect cannot be learned from the saccade-induced change of retinal inputs which have a latency of at least 50 ms to reach LIP/FEF and are suppressed by the saccade. Instead, the brain likely has access to only a stimulus’ pre-saccadic and post-saccadic retinotopic positions. For a stimulus flashed before the saccade, its post-saccadic position could be estimated according to the CD direction and strength. For a persistent stimulus, its post-saccadic position could additionally be provided by the reafference input after the saccade.

In this paper, we show that adding the continuity equation to the loss function helps neural network models learn the required connectivity patterns from just pre- and post-saccadic retinotopic positions of stimuli. The continuity equation is a hard constraint on locally conserved quantities such as electric charge. Neural activities, by contrast, can appear and disappear and are not conserved in general. However, in the special case of updating or controlling sustained neural population activities as representations or memories of variables, the activities are approximately conserved and the continuity equation can be a soft constraint as a regularizer in the loss function. We also studied whether the CD strength as a function of time, which controls the updating/remapping process, can be learned together with the connections (both randomly initialized) when we only specify the CD time window around the saccade. Finally, we examined whether the continuity loss enables learning when the time courses of stimuli’s retinotopic positions are heavily down sampled.

## 2 Results

We modeled transsaccadic updating in neural networks consisting of 100 units covering 100 deg of visual space along the saccade axis (Fig. 1a). The units were recurrently connected to each other with three sets of connections; two sets were gated by the CD signals for the two opposite directions of saccades and the third set was not gated. The units received feedforward, retinotopic inputs and their activities served as the model output. When the ungated connections follow a symmetric, center-excitation/surround-inhibition pattern (red curve in Fig. 1b) and the CD-gated connections follow an anti-symmetric, directional pattern (blue curve in Fig. 1b for rightward saccade/leftward updating; a mirror curve, not shown, for the leftward saccade/rightward updating), the network does transsaccadic updating of population activities representing stimuli’s retinotopic positions. Intuitively, the symmetric connectivity sustains the population activity bump evoked by a stimulus at a given retinotopic position and the CD-gated, anti-symmetric connectivity shifts the activity bump backward across the saccade. Because of the backward shift of the activity bump, a given cell shows forward RF remapping (Wang et al., 2025).

In this study, we initialized all the recurrent connections with random values and focused on the question of whether the required connectivity patterns can be learned under various conditions. We showed previously that the required connectivity patterns can be learned by using the desired output at every simulated time step as the teaching signal (Wang et al., 2024). We refer to this as the *full-output* condition (Fig. 1c). As we noted in Introduction, this condition may not be biologically plausible. We therefore considered the *partial-output* condition in which the desired output for only the pre- and post-saccadic positions of stimuli was provided for training and the intermediate, transitional trajectory was not specified.

### 2.1 Adding the continuity equation to the loss as a regularizer improves training results

We first reproduced our previous results (Fig. 2b, g, full-output condition). To match the remapping time courses in LIP and FEF, we used sluggish CD signals that started about 100 ms before the saccade onset and ended 100 ms after the saccade offset in the simulations. Since the exact form of the CD is unknown, we considered two different CD time courses, a Gaussian function and a step-like function (Fig. 2a, f); they produced similar results (Fig. 2b, g). For each saccade direction, we considered saccades of different magnitudes and the CD signal was multiplied by the magnitudes. The connectivity patterns after the training were similar to those in Fig. 1b. The actual outputs were similar to the desired outputs used for training although the former were not as smooth as the latter particularly during the transition from the pre- to post-saccadic retinotopic positions. Throughout the paper, the actual output plots show responses to inputs at test positions that are not included in the training. We show examples of rightward saccades but we obtained near identical results for leftward saccades.

**Fig. 2.**
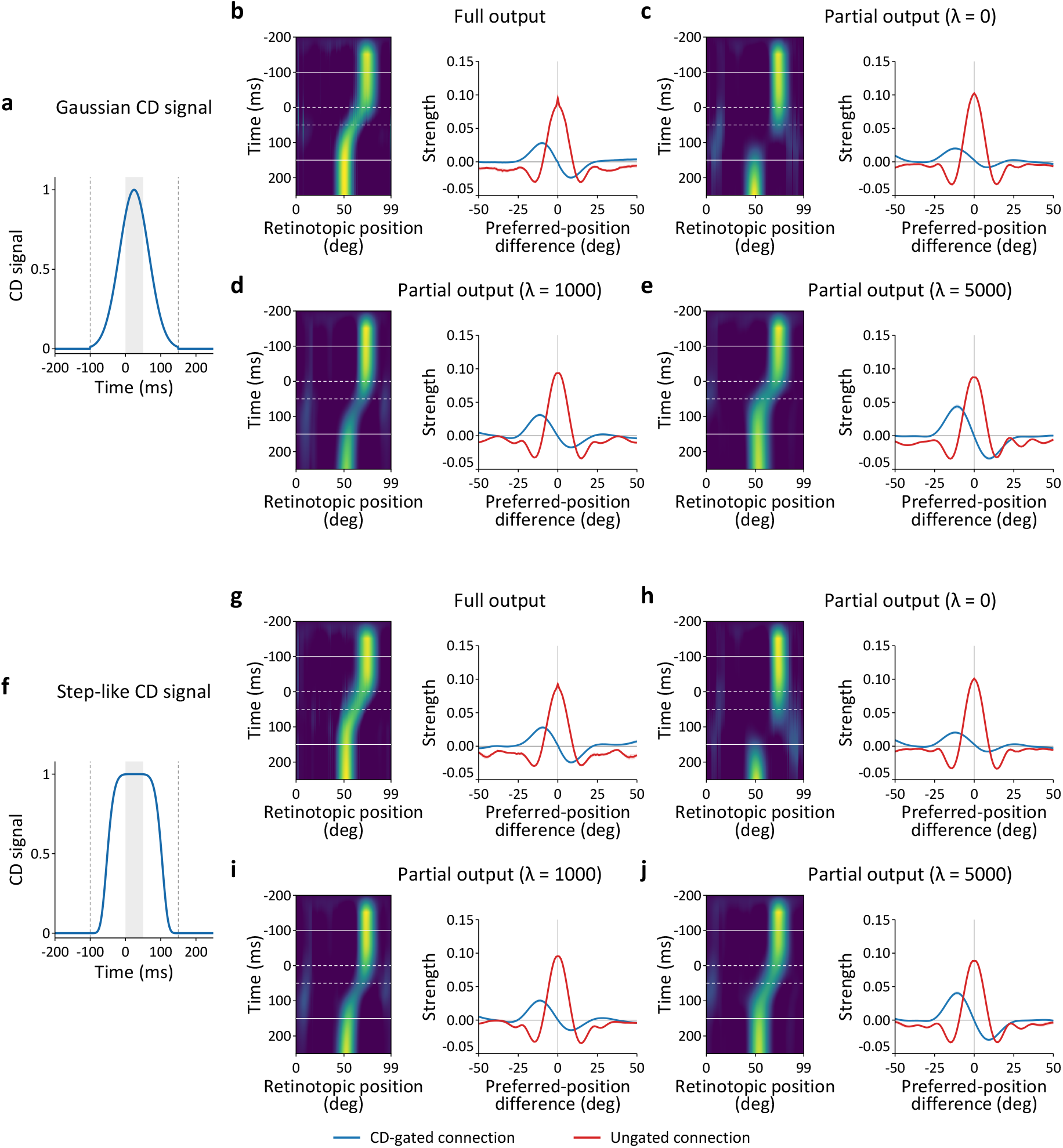
Actual outputs and recurrent connections after training under fixed Gaussian and step-like CD signals. (a) and (f) show Gaussian and step-like CD time courses, respectively; vertical dashed lines mark the CD time window and the shading marks the saccade time window. Under the Gaussian CD, (b) shows the results of the full-output condition, and (c)–(e) show the results of the partial-output condition with *λ* = 0, *λ* = 1000, and *λ* = 5000, respectively. (g)–(j) show the corresponding results under the step-like CD signal. The example heatmaps show the actual output of units tuned to different retinotopic positions (horizontal axis) as a function of time (vertical axis). Solid and dashed horizontal lines in the heatmaps mark the CD window and saccade window, respectively. The blue and red curves show the mean CD-gated and ungated recurrent connections, respectively, as a function of the difference between connected units’ preferred retinotopic positions (RF centers). Only the CD-gated connections for rightward saccades are shown. The shaded areas (often too small to be visible) indicate 1 SEM across connection weights with the same preferred-position difference.

Next, we applied the same training procedures to the partial-output condition. The results (Fig. 2c, h, *λ* = 0 to be defined below) were clearly not as good as those for the full-output condition. The CD-gated asymmetric connections were not fully developed. More importantly, the actual output trajectory stayed at the pre-saccadic position well after the CD signal started, and then jumped from the pre- to post-saccadic positions, contradicting the gradual increase of forward remapping magnitude observed in LIP and FEF (Wang et al., 2024).

Since the gradual increase of forward remapping magnitude implies a gradual backward updating of population activities (Qian et al., 2023; Wang et al., 2025), it seems reasonable to encourage the continuity of the population activities. We therefore added the continuity equation as a term in the loss function, with a weighting factor *λ* (Method 4.2).

Figs. 2c-e and 2h-j show the simulation results for the Gaussian and step-like CD signals, respectively, which are similar to each other. As the continuity loss becomes more important (*λ* increases from 0 to 5000), the output of the trained model becomes more continuous in time, as expected, and interestingly, the CD-gated connections become closer to anti-symmetric, a pattern known to preserve the shape of the activity bump during updating (Zhang, 1996).

### 2.2 The continuity regularization improves learning from down-sampled outputs

The above simulations were motivated by biological plausibility. We next examined whether the continuity loss could be useful in engineering applications. In many training situations, the target output may be available only at a subset of time points rather than at every step. We therefore asked whether the continuity regularization could help learning when the target output was limited in time. We started by down-sampling the desired output trajectory every 10, 20, 50 ms, and found that in each case, the training with the continuity loss was successful. We thus tried the extreme case of training with only two time samples, one at the pre-saccadic position and the other at the post-saccadic position (Fig. 3a). We refer to this as the *two-sample-output* condition. The results in Fig. 3b show that without the continuity loss (*λ* = 0), the population activity almost disappeared between the two supervised time points and did not form a clear moving bump. The CD-gated connectivity pattern was almost flat at zero and the ungated connectivity pattern was not fully developed. With the continuity loss (*λ* = 1000 and 5000), the activity bump was clearly maintained and shifted to the post-saccadic position. The CD-gated connections became more anti-symmetric, while the ungated connections showed a more symmetric, center-excitation and surround-inhibition pattern (Fig. 3c, d).

**Fig. 3.**
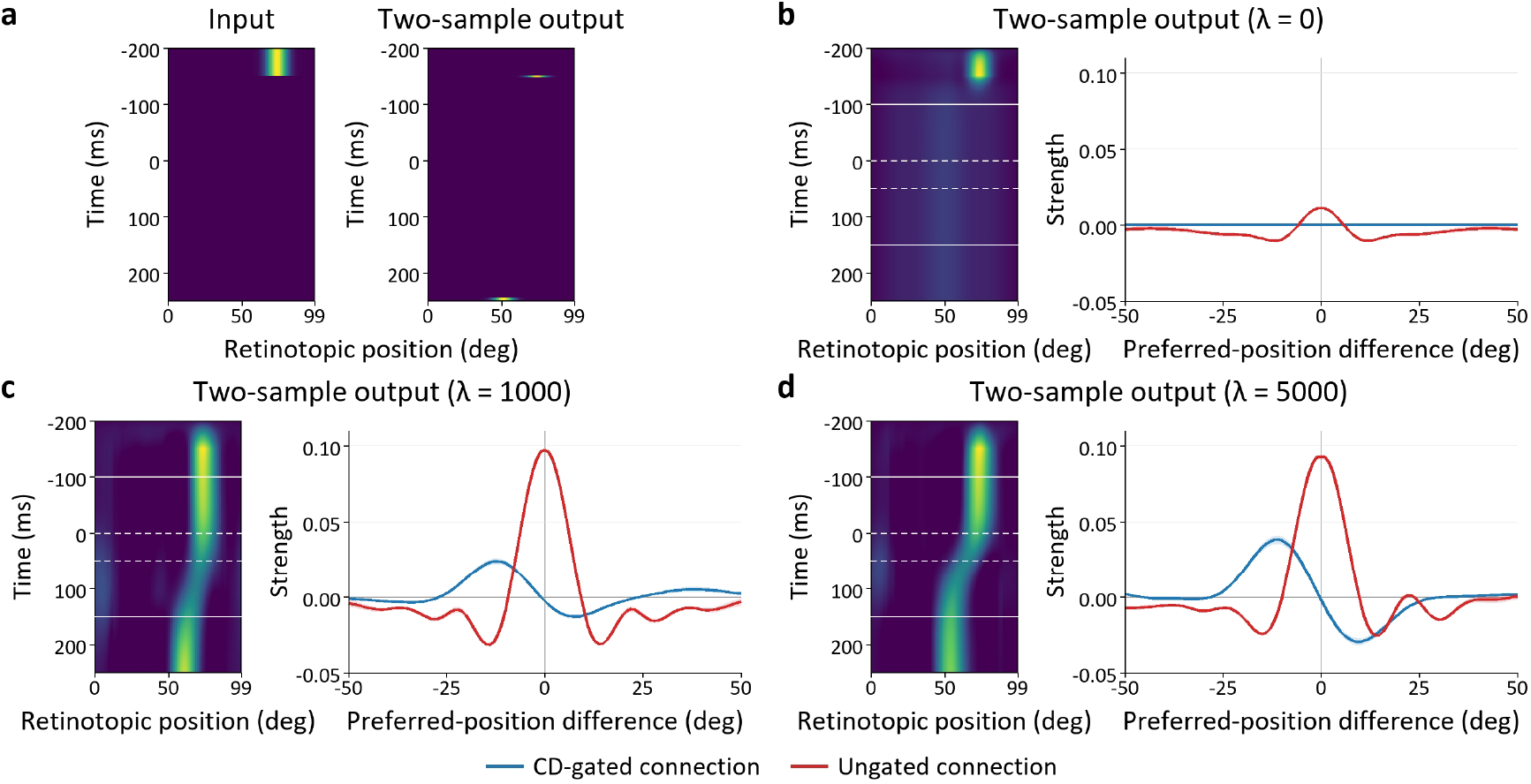
Actual outputs and recurrent connections after training under the two-sample-output condition with a fixed Gaussian CD signal. (a) An example of a brief visual input and the desired output provided only at *−*150 and 249 ms relative to saccade onset at 0 ms. The activities at remaining time steps were not specified. The target at 249 ms is broadened for visibility; supervised training was applied only at the two specified time steps. (b)–(d) show the actual outputs and mean recurrent connections for *λ* = 0, *λ* = 1000, and *λ* = 5000, respectively. The presentation formats are identical to those for the heatmaps and connections in Fig. 2.

### 2.3 The continuity regularization improves learning of the CD control signal

In the simulations above, we fixed the CD time course to be either Gaussian or step-like and trained neural networks to learn the connectivity patterns. We next examined whether the continuity loss could help with learning the CD signal, which controls the updating process, together with the connectivity patterns. Biologically, the CD signal derives from a copy of the saccade motor command but they may not have the same form as functions of time. We therefore only specified the time window for the CD signal but randomly initialized its value at each time step, similar to the random initialization of all the recurrent connections.

Under the full-output condition, the actual output after learning closely followed the desired updating trajectory (Fig. 4a). The learned CD signal was negative throughout its time course (sum = *−*33.56, see Method 4.4), whereas the CD-gated connections were near zero and flipped the sign. This combination works because the updating process depends on the product of the CD signal and the CD-gated connections. However, such an uneven division of magnitudes may not be robust in biological or engineering implementations.

**Fig. 4.**
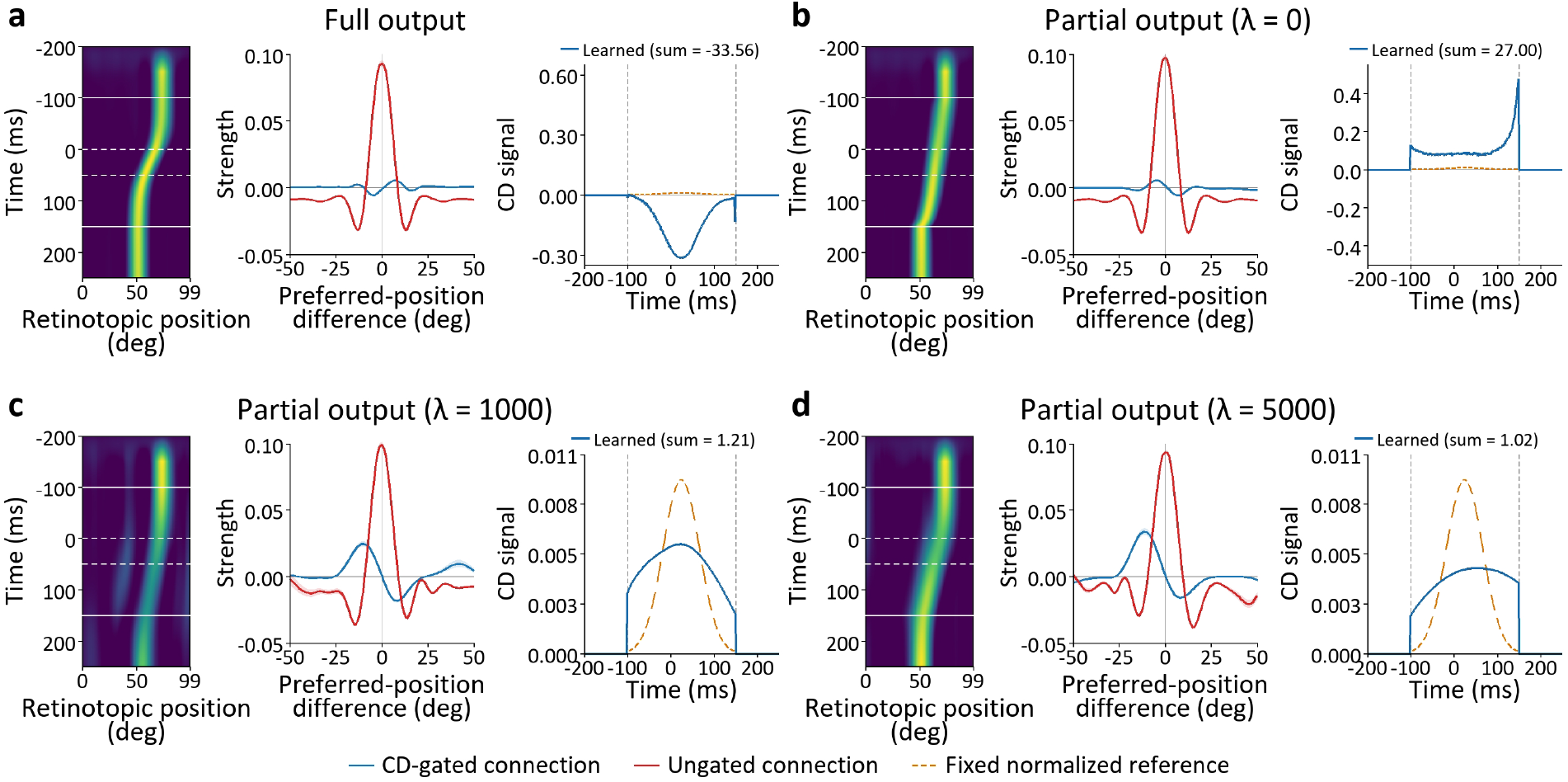
Actual outputs, recurrent connections and CD control signals after training. (a) The full-output condition, and (b)–(d) the partial-output condition with *λ* = 0, *λ* = 1000, and *λ* = 5000, respectively. In each panel, the left and middle plots show the actual output and connectivity patterns with the same format as those for Fig. 2. The right plot shows the learned CD time course (blue) and the normalized fixed Gaussian CD time course (orange dashed) as a reference; the sum of the learned CD time course is reported above each plot. Vertical dashed lines in the CD plots mark the CD window.

We then simulated learning under the partial-output condition. Without the continuity regularization, the actual output after learning also followed a continuous trajectory (Fig. 4b). Here, the learned CD signal had a large amplitude (sum = 27.00) with a spike at the end, and the CD-gated connections were small. Again, this solution works because the updating process depends on the product of the CD signal and the CD-gated connections but may not be robust.

With the continuity loss included (*λ >* 0, Fig. 4c, d), the learnable CD parameters influence the loss both directly via the continuity loss and indirectly via their gating of some recurrent connections (see Methods). Because of the form of the continuity equation used in our simulations, the CD signal is biased to be positive and to sum to 1 (Eq. (18)). With *λ* = 1000 and *λ* = 5000, the learned CD time courses were indeed positive and summed to 1.21 and 1.02, respectively. The magnitudes of the CD signal and the CD-gated connections were more evenly divided. Thus, the continuity loss may help improve the robustness of the learned solutions.

## 3 Discussion

We studied how to train neural network models to learn continuous updating and control of neural population activities over time, using transsaccadic processing as an example. To learn the connectivity patterns needed for continuous updating, we previously used desired output trajectories sampled at every time step as the teaching signal. Here we found that by adding the continuity equation to the loss function, we can greatly reduce the number of time samples needed for training. Specifically, we successfully trained the model with only the pre- and post-saccadic intervals without providing intermediate samples for the transitions. This setup is consistent with the neurobiological argument that before the brain learns to perform transsaccadic updating, it may not have access to the transitions. The continuity loss also helps with successful training when the desired output was reduced further to one pre-saccadic and one post-saccadic time point. This could be useful for engineering applications because dense time samples for training may be costly to obtain. Finally, we showed that the CD signal, which controls the updating process, can be learned together with the connectivity patterns, and the continuity loss may improve the robustness of the learned solution.

At the output of a network, a variable of interest can be represented either as a population activity of many cells (distributed representation) or as a monotonic function of a single cell’s activity (single-cell representation). Both types of representations have been used in neural modeling (Cueva, Ardalan, Tsodyks, & Qian, 2021; Sussillo & Abbott, 2009; Wang et al., 2024; Zipser & Andersen, 1988). The continuity loss can be easily applied to distributed representations but not single-cell representations. Consider, for example, these two types of representation for a joint angle changing from 10 to 20 degrees. In the single-cell representation, the cell’s activity increases with the angle so that the activity is far from conserved and the continuity equation is not applicable. By contrast, in a distributed representation, the angle is represented by the population activity bump among the cells tuned to different angles. The change of the angle is a shift of this activity bump as a wave and during the shift, the total activity is approximately conserved. Then, it is natural to add the continuity equation as a soft constraint in the loss function. This suggests an advantage of distributed representations over single-cell representations although the former require more cells than the latter. This advantage is different from (but not mutually exclusive with) the traditional view that distributed representations are more robust than single-cell representations. Distributed representations and activity waves are ubiquitous in the brain (Muller, Chavane, Reynolds, & Sejnowski, 2018). Whether and how the brain might use the continuity loss with distributed representations to facilitate learning, and whether some cortical waves are related to the updating or control of neural activities, are open questions.

Although we focused on the problem of transsaccadic updating/control of stimuli’s retinotopic positions, we believe that the benefits of including a continuity loss in training neural networks may be applicable to many problems that involve updating or control of sustained neural activities over a period of time. They may include the updating of memory activities for sensory variables or ongoing control of motor-related variables such as joint angles. It would be interesting to explore such applications in future studies.

## Acknowledgement

This work was supported by US NIH/NEI (R01 EY032938) and National Natural Science Foundation of China (32030045 and 32061143004).

## Author contributions

NQ and ZW designed the study and the model. ZW wrote the code, simulated the model, and produced the figures. ZW, NQ, and MZ wrote the manuscript.

## 4 Methods

### 4.1 Recurrent neural networks

We used recurrent neural networks adapted from our previous model of transsaccadic remapping/updating (Wang et al., 2025, 2024). Each network consisted of two layers: a visual input layer and a recurrent LIP/FEF layer whose activities served as the output. For simplicity, we considered only the horizontal dimension and horizontal saccades. Each layer contained 100 units representing 100^*°*^ of space. The feedforward connections were translationally invariant, implemented as a convolution with a five-unit kernel, unit stride, and symmetric zero padding to preserve the retinotopic representation.

The output units were fully recurrently connected through three sets of weights. Two sets were multiplicatively gated by CD signals for opposite saccade directions, whereas the third set was ungated. For each trial, the CD signal was supplied through the channel corresponding to the saccade direction, and the other channel was zero. The fixed and learnable CD time courses are described in Section 4.4.

The activities of the recurrent units were updated according to

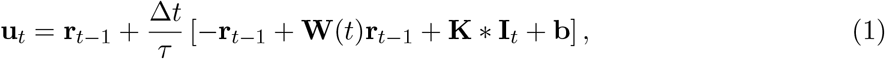

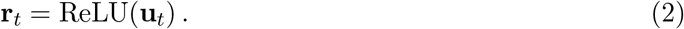

Here, **u**_*t*_ and **r**_*t*_ are the activation before rectification and firing rate of the recurrent units, respectively. The matrix **W**(*t*) specifies the recurrent connections, including the ungated and CD-gated components. The symbol *\** denotes spatial convolution, **K** is the feedforward kernel, **I**_*t*_ is the visual input, and **b** is the bias. We used the ReLU activation function, a time constant of *τ* = 10 ms, and a time step of Δ*t* = 1 ms. All recurrent activities were initially zero.

The recurrent weights were initialized independently from a uniform distribution over 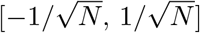, where *N* = 100, and the convolutional weights from a uniform distribution over 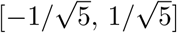. Both sets of weights were learned during training, without imposing symmetric or antisymmetric recurrent connectivity.

### 4.2 The loss function

We considered two types of loss for training. The first was the standard mean squared error between the desired (target) output and the actual output activities of the recurrent units. Let *T*_target_ denote the set of time steps at which a desired output *y*_*i,t*_ was provided. The loss was given by:

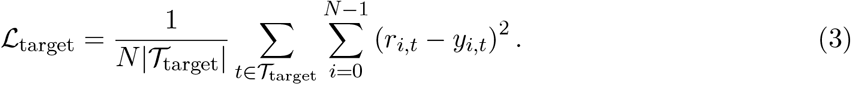

The average was over the |*T*_target_| time steps and the *N* = 100 spatial units. The condition-specific choices of *T*_target_ are specified below for the full-output, partial-output, and two-sample-output conditions. This was the only loss used in some of our simulations.

For the other simulations, we added a second loss according to the continuity equation for local conservation of a quantity *ρ*(*x, t*) moving with velocity *v*(*x, t*):

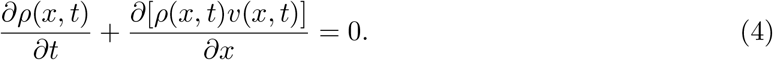

In our case, the quantity *ρ*(*x, t*) was the output activity bump representing the retinotopic position of a stimulus. When the velocity was spatially uniform, *v*(*x, t*) = *v*(*t*), Eq. (4) reduced to

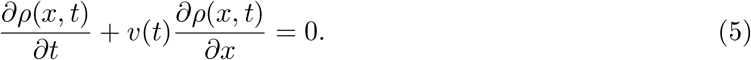

When the CD-gated recurrent connectivity is a spatial derivative of the ungated, symmetric connectivity, we know (Zhang, 1996):

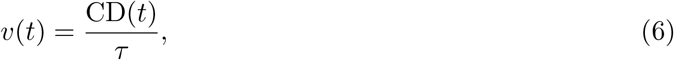

where CD(*t*) is the time course of the CD signal and *τ* is the network time constant. The total accumulated updating magnitude is given by:

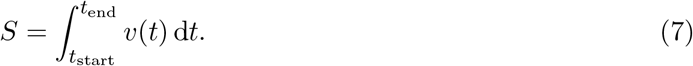

In our case, all the recurrent connections were initialized randomly, and we found that training was more stable when we used the following equivalent form of Eq. (6):

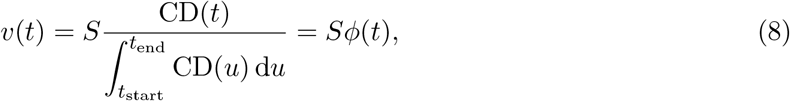

where *ϕ*(*t*) is defined as the normalized CD signal, and *S* is set to the desired updating/remapping magnitude.

For spatial unit *i* and time step *t*, the discretized continuity residual was

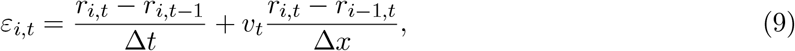

with Δ*t* = 1 ms and Δ*x* = 1^*°*^. Let *T*_cont_ denote the time steps at which the continuity residual was evaluated. The continuity loss was the mean squared residual over the |*T*_cont_| time steps and the *N −* 1 spatial differences,

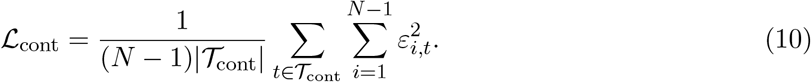

The total loss was

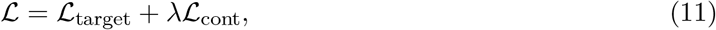

where *λ ≥* 0 controlled the relative importance of the two terms.

### 4.3 Transsaccadic updating of retinotopic positions

We trained our neural networks on the task of transsaccadic updating of stimuli’s retinotopic positions (Wang et al., 2024). Specifically, the output units should have the population activity pattern representing the updated retinotopic position of an input stimulus across saccades. We assumed that positional coding of the output units was fixed without changing with transient RF shifts (unaware decoder) (Qian et al., 2023; Wang et al., 2025). Both the input and desired output were Gaussian activity patterns with unit peak amplitude and a standard deviation of 5^*°*^. We considered brief input stimuli, with one stimulus per training trial. The stimuli appeared for 50 ms before saccades and then disappeared. After stimulus disappearance, the network was required to maintain the stimulus representation and use the CD signal to update its position in the direction opposite to the saccade, by an amount equal to the saccade amplitude.

We considered three training conditions. The full-output condition provided the desired output activities at each simulated time step. During the transition, the center of the desired output followed the cumulative normalized CD time course:

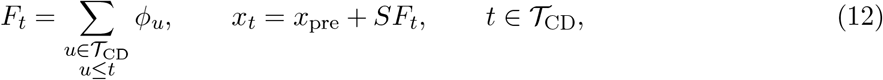

where *x*_pre_ denotes the pre-saccadic center. The partial-output condition specified the desired output activities for only the pre- and post-saccadic retinotopic positions of a stimulus. For our simulations with a total of 450 time steps, the activities for pre- and post-saccadic positions were specified for [50, 100) and [350, 450), respectively, corresponding to [*−*150, *−*100) and [150, 250) ms relative to saccade onset. For all the other time steps, the desired output was unspecified and no target error was generated. In the down-sampled-output condition, the desired output was specified every *d* model time steps, with *d* = 10, 20, or 50. As an extreme case, we also considered the two-sample-output condition, in which the desired output was specified only at model steps 50 and 449, corresponding to *−*150 and 249 ms relative to saccade onset. For simulations with *λ >* 0, *T*_cont_ consisted of model steps in [100, 351) for the partial-output condition and [50, 450) for the down-sampled-output condition.

### 4.4 Fixed and learnable CD signals for controlling updating/remapping

For some simulations, we fixed the CD signal to have either a Gaussian or a step-like time course, with an amplitude proportional to the updating/remapping magnitude (which, in turn, equaled the saccade amplitude):

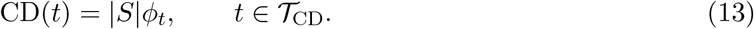

The CD signal was zero outside *T*_CD_.

The normalized time course for Gaussian CD was

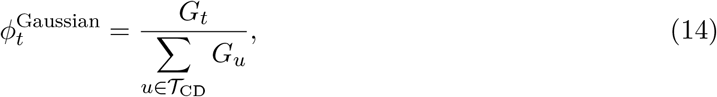

with

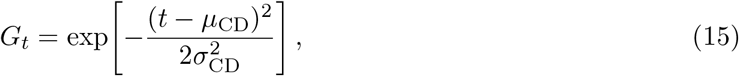

where *µ*_CD_ = 24.5 ms, and *σ*_CD_ = (250*/*6) ms relative to saccade onset at 0 ms.

The normalized time course for step-like CD was

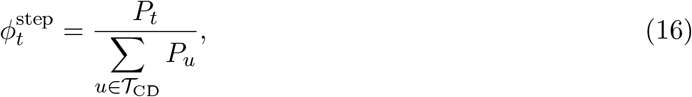

with

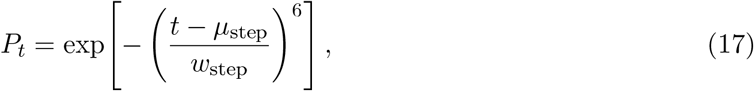

where *µ*_step_ = 24.5 ms and *w*_step_ = 81.6 ms, relative to saccade onset.

For the other simulations, we let CD(*t*) at time steps within *T*_CD_ be learnable parameters. Their values were initialized independently from a Gaussian distribution *N*(0, (1*/*250)^2^). In the continuity-regularized learnable-CD conditions, the velocity term was

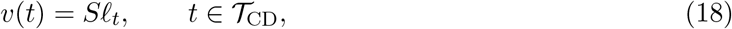

where *ℓ*_*t*_ denotes the learned CD time course. Given Δ*t* = 1 ms, Learning the required monotonic shifts of activity bumps favored positive values of *ℓ*_*t*_ with Σ_*t*_ *ℓ*_*t*_ close to 1. We trained neural networks to learn the connections and the CD signals together under both the partial-output and full-output conditions. For the full-output condition, the transitions in the desired trajectories were created with the Gaussian CD time course mentioned above, independent of the actual CD time courses during the training.

